# Dynamic interactions of lymphatic vessels at the hair follicle stem cell niche during hair regeneration

**DOI:** 10.1101/548768

**Authors:** Daniel Peña-Jimenez, Silvia Fontenete, Diego Megias, Coral Fustero-Torre, Osvaldo Graña-Castro, Donatello Castellana, Robert Loewe, Mirna Perez-Moreno

## Abstract

Lymphatic vessels (LV) are essential for skin fluid homeostasis and immune cell trafficking, but whether LV are associated with hair follicle (HF) regeneration is not known. Here, by using steady and live imaging approaches in mouse skin, we show that lymphatic capillaries distribute to the anterior permanent region of individual HF and interconnect neighboring HF at the level of the HF bulge, in a hair follicle stem cell (HFSC)-dependent manner. LV further connect individual HF in triads and dynamically flow across the skin. Interestingly, at the onset of the physiological HFSC activation, or upon pharmacological or genetic induction of HF growth, LV transiently expand their caliber suggesting an increased tissue drainage capacity. Interestingly, the physiological LV caliber increase is associated with a distinct gene expression correlated to ECM and cytoskeletal reorganization. Using mouse genetics, we show that the depletion of LV blocks the pharmacological induction of HF growth. Our findings define LV as components of the HFSC niche, coordinating HF connections at tissue-level, and provide insight into their functional contribution to HF regeneration.

## Introduction

Lymphatic vessels (LV) play fundamental homeostatic functions including the balanced transport of fluids and macromolecules, the local coordination of immune responses, as well as immune cell trafficking to regional lymph nodes (Skobe & Detmar, 2000). After years of scientific discovery, much has been learned about the distinctive characteristics of LV, including the molecular markers Prospero-related homeobox 1 (Prox1), Lymphatic vessel endothelial hyaluronan receptor 1, (LYVE-1) and podoplanin, as well as critical regulatory signals that govern their development and fundamental functions in tissues (Potente & Makinen, 2017; Wang & Oliver, 2010; Yang & Oliver, 2014; Zheng *et al.*, 2014).

In skin, LV are organized in structured polygonal patterns, consisting of one subcutaneous plexus, and a more superficial plexus located in the dermis, near the blood vessels (Braverman, 1989). Several studies have contributed to our understanding of the orderly organization of LV in the skin (Skobe & Detmar, 2000; Tripp *et al.*, 2008). These studies have provided insight into the existence of branches of lymphatic capillaries that extend to the HF and drain into the subcutaneous collecting LV, presumably through connections of blind capillaries with the dermal papillae (dp) (Forbes, 1938), a condensate of dermal fibroblasts that provides a specialized microenvironment (Millar, 2002; Sennett & Rendl, 2012; Yang & Cotsarelis, 2010). LV may also facilitate the entry of immune cells to the HF epithelium, a source of chemokines that regulate the trafficking of epidermal Langerhans cells and dermal dendritic cells (Nagao *et al.*, 2012), the distribution and differentiation of Langerhans cells (Wang *et al.*, 2012), and the tropism of skin resident memory T cells (Adachi *et al.*, 2015). However, despite the role of LV in facilitating immune cell trafficking to HF, less is known about their coordinated connections during HF cycling and functional implications.

In adult skin, HF exhibit a lifetime polarized pattern of growth and regeneration across the tissue modulated by stimulatory and inhibitory signals (Plikus *et al.*, 2011; Widelitz & Chuong, 2016). The cyclic regeneration of HF involves phases of growth (Anagen) via regression (Catagen) to relative quiescence (Telogen) (Geyfman *et al.*, 2015). The entry of resting HF into Anagen requires the activation of HFSC located in the HF bulge (Cotsarelis *et al.*, 1990; Tumbar *et al.*, 2004), and the expansion of their progenitors found in the secondary hair germ, giving rise to a new Anagen HF (Greco *et al.*, 2009; Rompolas *et al.*, 2012; Tumbar *et al.*, 2004). Anagen HF grow until other instructive signals promote their regression giving rise to a new HF cycle. In past decades, a wealth of knowledge has yielded valuable insight into the role of major stimulatory and inhibitory signals in governing the orchestrated activation of the HF cycle (Blanpain & Fuchs, 2009; Lee & Tumbar, 2012; Plikus & Chuong, 2014), including local self-activation signals (Hsu *et al.*, 2011), the contribution of other cells in the tissue macroenvironment (Ali *et al.*, 2017; Brownell *et al.*, 2011; Castellana *et al.*, 2014; Festa *et al.*, 2011; Rivera-Gonzalez *et al.*, 2016), as well as long-range signaling waves across the skin (Plikus *et al.*, 2011).

Blood vessels have also been found associated around HF exhibiting a coordinated dynamic reorganization during HF cycling (Mecklenburg *et al.*, 2000; Yano *et al.*, 2001). In addition, HFSC closely associate to a venule annulus (Xiao *et al.*, 2013). In unperturbed skin, the occurrence of angiogenesis, the growth of new capillaries from pre-existing blood vessels, has been observed during Anagen (Mecklenburg *et al.*, 2000). The epidermal expression of the Vascular Endothelial Growth Factor A (VEGF-A) (Detmar, 1996) induces perifollicular angiogenesis and sustains HF growth; conversely, inhibition of VEGFA leads to a delay in HF growth accompanied by reductions in HF size and perifollicular vascularization (Yano *et al.*, 2001). Overall, these results exposed that HF and blood vessels form a functional operative system. In contrast, less is known about the role of LV in regulating this process. Here, we show that lymphatic capillaries are novel components of the HFSC niche, coordinating HF connections at tissue-level and provide insight into their functional association to the HF cycle.

## Results

### Lymphatic capillaries distribute in the vicinity of HFSC in a polarized manner

To investigate the association between LV and HFSC, we first defined the lymphatic distribution at HF in mouse backskin. To this end, we performed immunofluorescence analyses using antibodies to the lymphatic marker LYVE-1 as well as to alpha smooth muscle actin (αSMA), enriched in the arrector pili muscle (apm). Lymphatic capillaries distributed in a polarized manner, aligned to the anterior side of the HF, opposite to the distribution described for the apm (Fujiwara *et al.*, 2011), ascending as blind capillaries along the HF permanent area towards the epidermis until the infundibulum (Fig 1A). This distribution was different from the one observed in the ear skin, which presented parallel lymphatic capillaries that were not associated with HF (Fig EV1A). In the backskin, lymphatic capillaries densely distributed to the anterior side of HF at Tenascin C areas (Fig 1B), a glycoprotein of the extracellular matrix (ECM) enriched in the HF bulge (Tumbar *et al.*, 2004). To interrogate the existence of a lymphatic association with both embryo and adult HFSC, we next determined the lymphatic distance to Lhx2^+^ SC (Fig 1C and D) (Rhee *et al.*, 2006) and CD34^+^ HFSC (Fig 1E and F) (Blanpain *et al.*, 2004), respectively. Immunofluorescence analyses of backskin sections of P5 and P12 mice (Fig 1C and D) and P49 mice (Fig 1E and F) revealed that lymphatic capillaries distributed to the proximity of HFSC, within a distance inferior of 3 μm, within the ratio expected for components of the HFSC niche (Beck *et al.*, 2011).

**Figure 1.**
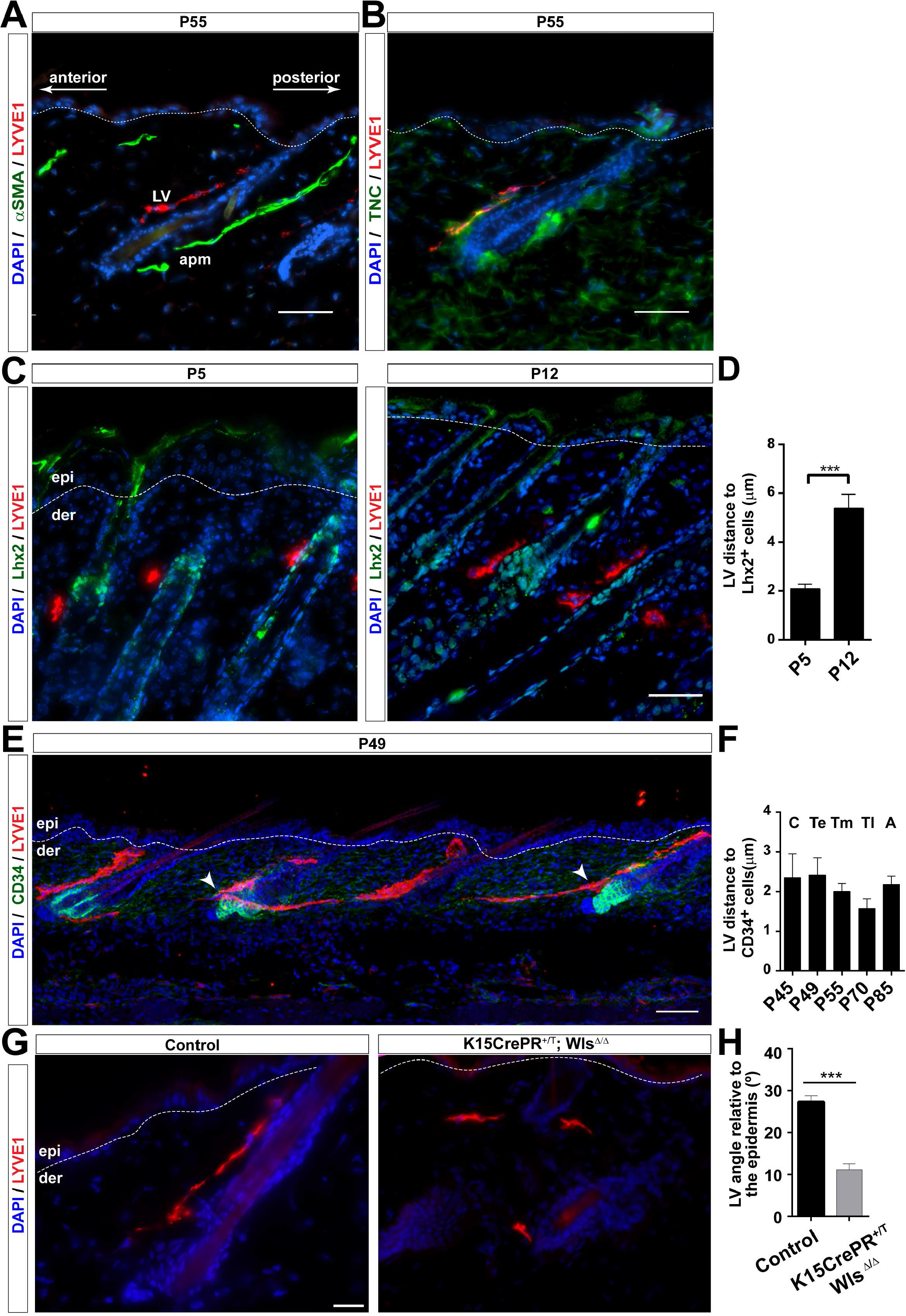
LV interact with HF in a functional HFSC niche dependent manner. A. Adult mouse backskin sections immunostained for LYVE1 (red), αSMA (green) and counterstained with DAPI (blue). n= 3 – 4 skin samples per mouse, n= 3 - 4 mice. Bar, 50 μm. LV, lymphatic vessels; apm, arrector pili muscle. B. Backskin sections of P49 mouse skin immunostained for LYVE1 (red), Tenascin-C (green) and counterstained with DAPI (blue). n= 3 – 4 skin samples per mouse, n= 3 - 4 mice. Bar, 50 μm. epi, epidermis; der, dermis; TNC, Tenascin-C. C. Adult mouse backskin sections from different postnatal (P) days immunostained for LYVE1 (red), Lhx2 (green) and counterstained with DAPI (blue). n= 3 – 4 skin samples per mouse, n= 3 - 4 mice. Bar, 50 μm. epi, epidermis; der, dermis. D. Histogram of the quantification of the LV distance to HFSC positive for Lhx2. Data represent the mean value ± SEM. *p< 0.05; **p < 0.01; ***p < 0.001. E. Backskin sections of P49 mouse skin immunostained for LYVE1 (red), CD34 (green) and counterstained with DAPI (blue). n= 3 – 4 skin samples per mouse, n= 3 - 4 mice. Bar, 100 μm. epi, epidermis; der, dermis. F. Histogram of the quantification of the LV distance to HFSC positive for CD34. n= 3 – 4 skin samples per mouse, n= 3 - 4 mice. Data represent the mean value ± SEM. *p< 0.05; **p < 0.01; ***p < 0.001. G. Backskin sections of K15CrePR^+/T^; Wls^Δ/Δ^ and Control K15CrePR^+/+^; Wls^flox/flox^ mice immunostained for LYVE1 (red), and counterstained with DAPI (blue). n= 3 – 4 skin samples per mouse, n= 3 - 4 mice. Bar, 100 μm. epi, epidermis; der, dermis. H. Histogram of the quantification of the LV angle relative to the epidermis. n= 3 – 4 skin samples per mouse, n= 3 - 4 mice. Data represent the mean value ± SEM. *p< 0.05; **p < 0.01; ***p < 0.001.

Next, we explored if functional HFSC create a niche promoting the continuous association of LV with the HF bulge. Wnt signaling features prominently in HFSC and regulates their properties and functional activity (Choi *et al.*, 2013). Thus, we reduced the expression of Wnt ligands in HFSC by ablating the expression of Wls in the Keratin 15 (K15) HFSC compartment, using the K15CrePR^+/T^; Wls^Δ/Δ^ conditional mouse model (Choi *et al.*, 2013; Myung *et al.*, 2013). Wntless (Wls) binds to Wnt ligands and controls their sorting and secretion (Carpenter *et al.*, 2010). Under these conditions, HFSC remain quiescent, exhibiting a reduction in proliferation, but HF are largely maintained (Choi *et al.*, 2013; Myung *et al.*, 2013). Consistent with those findings, HF remained present in the skin (Fig 1G) despite the loss of Wls, as confirmed by RT-qPCR analyses of CD34+ α6 integrin+ FACS-isolated cells (Fig EV1B). LYVE1 immunofluorescence analyses revealed that the organized association of LV with HF was disrupted, and LV were found distributed parallel to the epidermis and distant from HF bulge areas (Figure 1G and H), reminiscent to the organization of LV in the ear skin (Fig EV1A). Overall, these results indicate that HFSC create a niche for lymphatic endothelial cells, and sustain a polarized pattern of LV, which in turn, interconnect neighboring HF at the level of the HF bulge across the skin.

### Lymphatic capillaries start associating with HF during morphogenesis

To gain further insight into the establishment of the LV – HF association, we next turned to HF embryogenesis. HFSC emerge in the early HF placode stage at E15.5, giving rise to the hair germ (E16.5), hair pegs (E17.5) and embryo anagen HF until the first postnatal HF cycle. Whole mount immunofluorescence analyses of LYVE-1 at E15.5 – E17.5 embryo stages (Fig 2A – C) revealed that from E15.5, when HF placodes were already visible, nascent networks of anastomosed LV start to form below areas of HF growth. HF start to develop via epithelial – mesenchymal inductive signals stemming from the dermal papilla (dp) (Sennett & Rendl, 2012); however, no apparent lymphatic association with the dp was observed at E15.5 or E16.5, and LV were rather aligned in parallel to the epidermis, representative of lymphatic collecting vessels (Fig 2A and B). Interestingly, at E17.5 upward grows of lymphatic capillaries started to branch out from collecting vessels towards the papillary dermis and distributed at HF sites (Fig 2C). Overall, these results indicate that the development of HF is coupled with the recruitment of lymphatic capillaries to HF sites, likely subsequent to HF specification.

**Figure 2.**
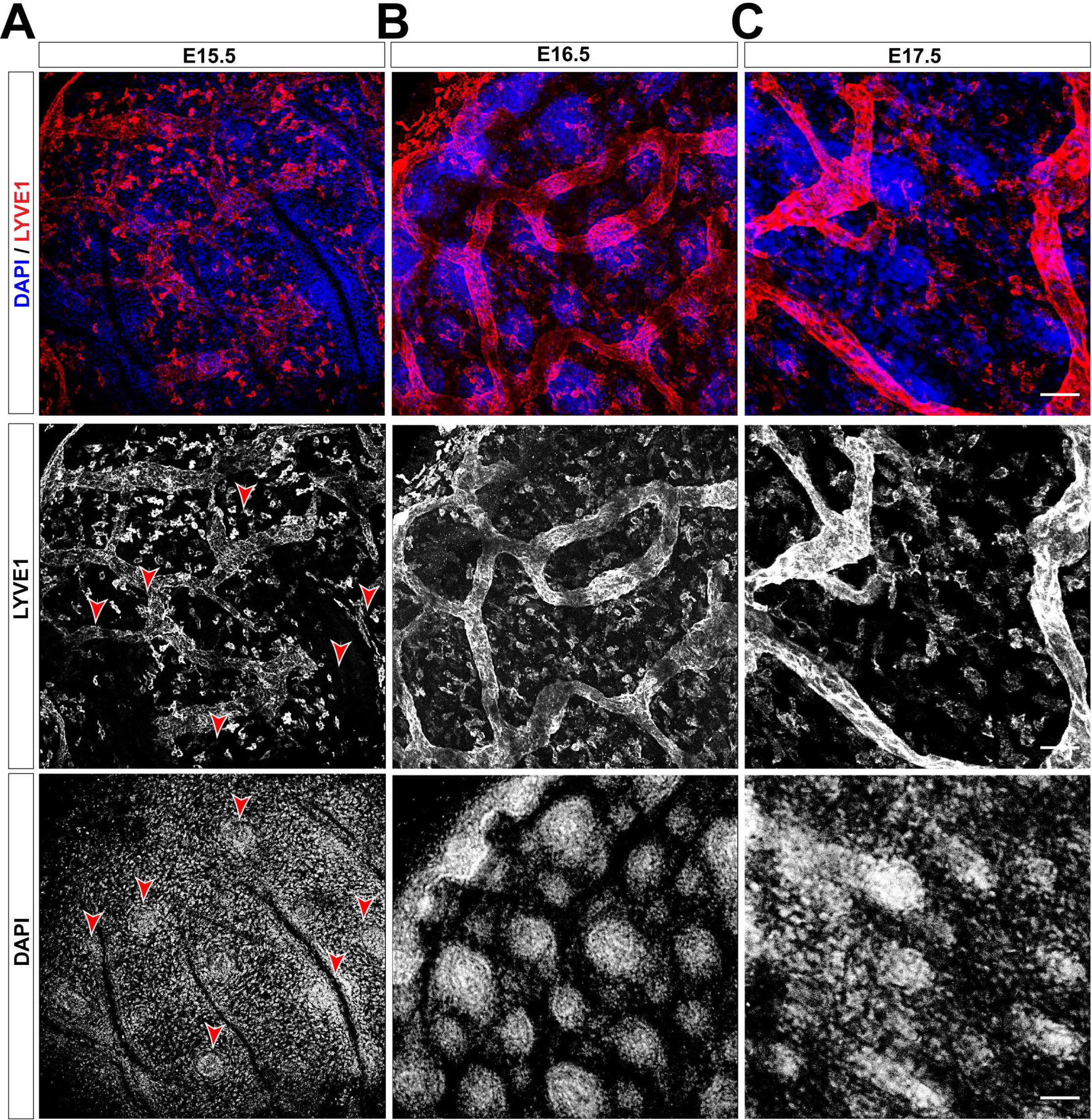
LV associate with HF during development. A. Distribution of LV and HF at the mouse embryo stage E15.5. Maximum projection images of whole mount immunofluorescence analyses using LYVE1 (red) as lymphatic endothelial marker and counterstained with DAPI (blue). Red arrowheads denote HF placodes. n= 3 – 4 embryos. Bar, 50 μm. B. Distribution of LV and HF at the mouse embryo stage E16.5. Maximum projection images of whole mount immunofluorescence analyses using LYVE1 (red) as lymphatic endothelial marker and counterstained with DAPI (blue). n= 3 – 4 embryos. Bar, 50 μm. C. Distribution of LV and HF at the mouse embryo stage E17.5. Maximum projection images of whole mount immunofluorescence analyses using LYVE1 (red) as lymphatic endothelial marker and counterstained with DAPI (blue). n= 3 – 4 embryos. Bar, 50 μm.

### Lymphatic vessels interconnect triads of HF across the backskin

To further analyze the organization of the lymphatic association with HF we conducted whole mount immunofluorescence analyses of LYVE-1 in adult skin sections (P70). This method allowed discerning an additional arrangement level, where individual LV-HF units associated further into triads (Fig 3A). The patterned polygonal organization of lymphatic – HF domains across the skin was more evident in 3D projection planes (Fig 3B and Movie EV1). Lymphatic capillaries were found associated along the permanent portion of individual HF. At the level of the HF bulge, lymphatic capillaries radiated and converged interconnecting three HF units. These units presented a common extending lymphatic capillary from each HF triad (Fig 3C and Movie EV2). The mechanisms involved in the lymphatic – HF patterning are interesting questions for the future, but globally, these results uncharted the existence of coordinated lymphatic arrays surrounding and interconnecting triads of HF across the skin.

**Figure 3.**
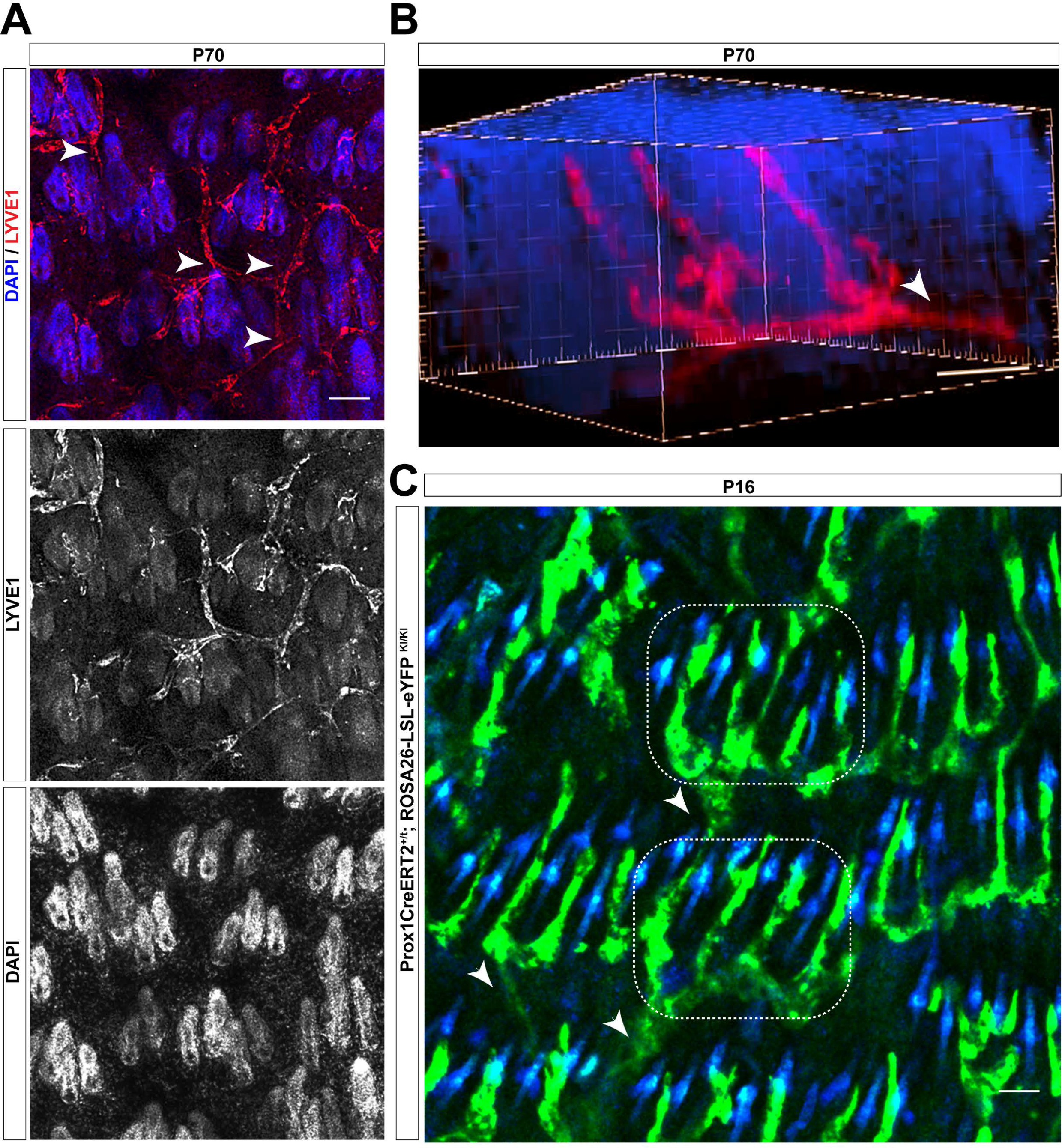
LV association with HF during the postnatal HF cycle. A. LV associated with individual HF further organize associating HF triads, which connect with other HF triads across the skin. P70, 70 d old mice. n= 3 - 4 mice. White arrowheads denote capillaries stemming from HF triads. Bar, 100 μm. B. 3D reconstructions of whole skin mounts showing a triad of HF connected by LV. n= 3 - 4 mice. Bar, 50 μm. C. Images of intravital microscopy analyses showing aligned HF rows in the backskin interconnected by LV in the Prox1CreERT2; Rosa-LSL-eYPF mice. Dotted boxes denote HF triads; White arrowheads denote capillaries stemming from HF triads interconnecting to other HF triads in adjacent rows. n= 3 – 4 mice. Bar, 50 μm.

### Lymphatic vessels continuously associate to the HF bulge during the HF cycle

We next explored if LV change their distribution to the HF bulge areas during the postnatal HF cycle. These analyses were performed in isolated backskin sections from matched skin areas at defined stages of the HF cycle, including the postnatal HF morphogenesis (Postnatal days 5 – 16, P5 – P16), and the first (P23 – P45) and the second (P45 – P69) HF cycle (Fig EV2) (Muller-Rover *et al.*, 2001). The latter exhibits a more extended Telogen that lasts for 3 – 4 weeks; therefore, to perform our comparative analyses we subdivided the second telogen into early telogen (Te, P49), mid telogen (Tm, P55), late telogen (Tl, P69), and included an Anagen stage (AVI, P85) (Muller-Rover *et al.*, 2001). These phases corresponded to the refractory and competent Telogen phases as previously documented (Plikus *et al.*, 2008).

Interestingly, at all HF stages, LV remained distributed in a polarized manner, positioned at the anterior side of the permanent region of the HF opposite to the apm, interconnecting neighboring HF at the level of the HF bulge (Fig EV2A). The greater lymphatic density was localized at the HF permanent region, exhibiting a significantly higher area at postnatal HF morphogenesis stages (P5-P16) (Fig EV2B). Consistent with this distribution, the relative LV/HF length increased from late Anagen stages to Telogen, when HF mostly consist of the permanent region (Fig EV2C). Conversely, this relative length was reduced during the transition from Telogen to Anagen (Fig EV2C).

These studies exposed that LV establish a continuous connection between neighboring HF at the level of the HF bulge across the skin throughout all phases of the HF cycle.

### Lymphatic vessels dynamically flow across triads of HF in the backskin

To further pursue the existence of a dynamic lymphatic flow between neighboring HF triads across the skin, we analyzed the lymphatic network by intravital microscopy, using a Prox1-CreERT2; ROSA26-LSL-eYFP reporter mouse. This transgenic mouse line expresses the tamoxifen-inducible Cre recombinase (CreER) under the control of the Prox1 gene promoter (Bazigou *et al.*, 2011), under a Rosa26-LSL-eYPP background. Prox1 is important for LV development, and it is expressed throughout life providing lymphatic identity (Hong *et al.*, 2002; Wigle & Oliver, 1999). These powerful tools allowed us to explore the dynamic association of lymphatic capillary networks with HF in living skin tissue. Our intravital microscopy analyses fully evidenced the continuum-patterned organization of lymphatic networks around HF in the backskin (Fig 3C). These analyses further allowed the visualization of triads of HF interconnected by lymphatic capillaries, aligned in parallel rows with an anterior to posterior disposition. Strikingly, the HF triads in each row interconnected with neighboring parallel rows, mainly through a LV stemming from one HF triad unit (Fig 3C), consistent with our prior observations (Fig 3B and Movie EV2).

We next assessed if LV dynamically streamed into the continuum-patterned lymphatic networks around HF in the backskin. To this end, TRITC – Dextran was administered in the mouse backskin followed by intravital microscopy. These results allowed the visualization of a continuous lymphatic flow interconnecting adjacent HF rows (Movie EV3).

These results exposed the existence of a dynamic HF communication through lymphatic vascularization, which potentially facilitates the spreading of signaling waves and immune cell trafficking across HF in the backskin.

### Lymphatic Endothelial Cells transiently increase their caliber at the onset of HF stem cell activation

Our previous findings raised the possibility that LV undergo dynamic flow changes at different stages of the HF cycle. We investigated this aspect by measuring the LV caliber in backskin sections during the first and the second HF cycle, conducting LYVE-1 immunofluorescence analyses. Intriguingly, the LV caliber was more pronounced at the onset of the Telogen to Anagen transition (Fig 4A and B). We next inspected more closely the LV morphology and observed that at late stages of Telogen, LV appeared more fenestrated displaying membrane protrusions compared to the more continuous and tight capillaries observed during Anagen. This structural variation, exhibiting a wide and irregular lumen, presumably accounts for regional differences in capillary permeability (O’Driscoll, 1992) influencing vascular exchange and increase tissue drainage capacity (Aebischer *et al.*, 2014; Betterman & Harvey, 2016).

**Figure 4.**
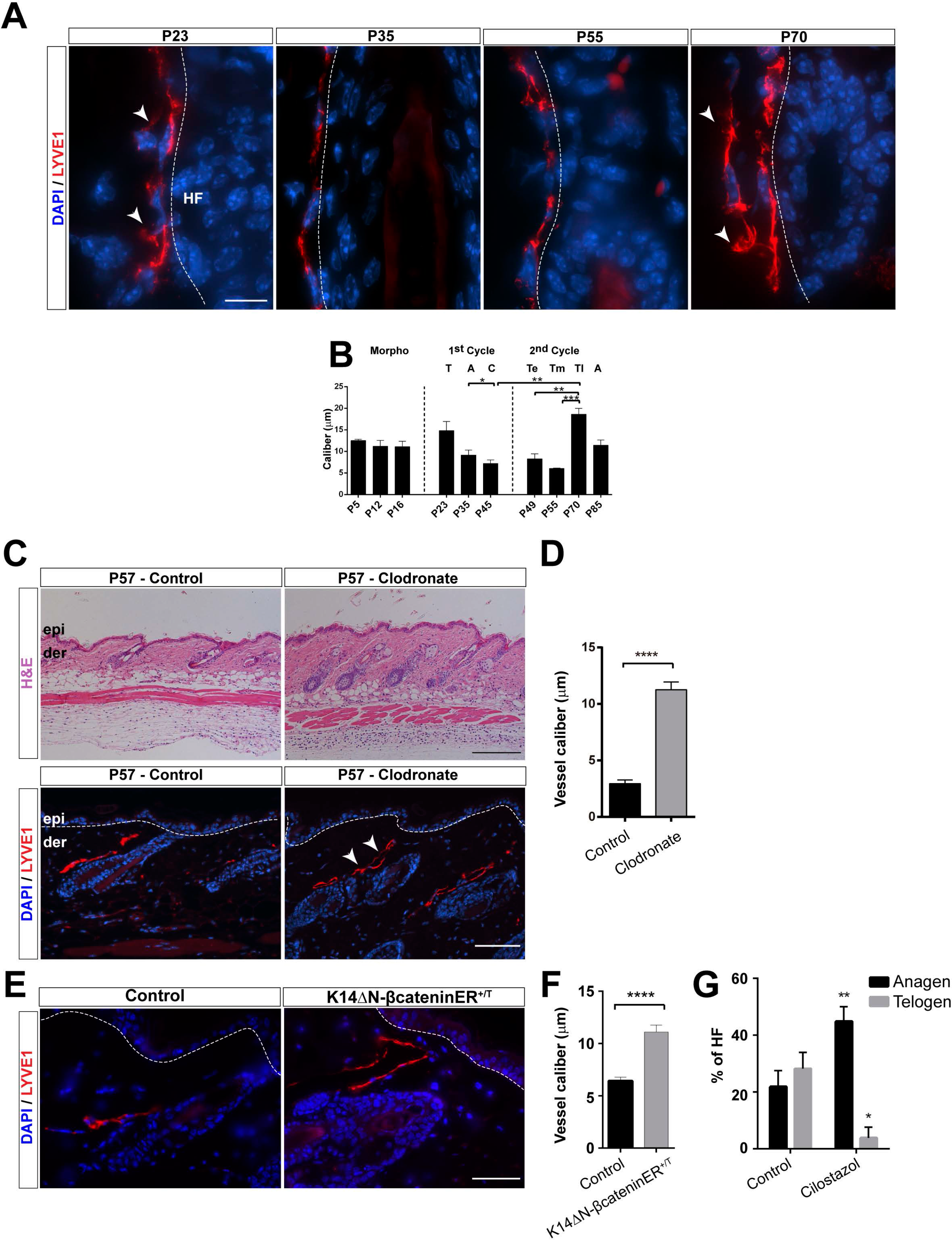
Dynamic reorganization of LV during the HF cycle. A. Adult backskin sections from different postnatal (P) days immunostained for LYVE1 (red) and counterstained with DAPI (blue). n= 3 – 4 skin samples per mouse, n= 3 - 4 mice. Bar, 10 μm. White arrowheads denote LV membrane protrusion and fenestrated areas. LV, lymphatic vessels; HF, hair follicle. B. Histogram of the caliber (μm) of LV at different postnatal days. n= 3 – 4 skin samples per mouse, n= 3 - 4 mice. A, Anagen; C, Catagen; T, Telogen; Te, early Telogen; Tm, mid Telogen; Tl, late Telogen. The data shown represent the mean value ± SEM. *p< 0.05; **p < 0.01; ***p < 0.001. C. Backskin sections from mice treated intradermally with Control or Clodronate liposomes, immunostained for LYVE1 (red) and counterstained with DAPI (blue). n= 3 – 4 skin samples per mouse, n= 3 - 4 mice. Bar, 200 and 100 μm. White arrowheads denote LV membrane protrusion and fenestrated areas. epi, epidermis; der, dermis. D. Histogram of the caliber (μm) of LV in the backskin of mice treated with Control or Clodronate liposomes. n= 3 – 4 skin samples per mouse, n= 3 - 4 mice. The data shown represent the mean value ± SEM. *p< 0.05; **p < 0.01; ***p < 0.001. E. Backskin sections of Controls and K14∆Nβ-cateninER^+/T^ mice immunostained for LYVE1 (red), and counterstained with DAPI (blue). n= 3 – 4 skin samples per mouse, n= 3 - 4 mice. Bar, 50 μm. epi, epidermis; der, dermis. F. Histogram of the LV caliber in skin sections from Controls and K14∆Nβ-cateninER^+/T^ mice. n= 3 – 4 skin samples per mouse, n= 3 - 4 mice. Data represent the mean value ± SEM. ***p < 0.001. G. Histogram of the percentage of HF in Telogen and Anagen present in the skin of control mice or mice treated with Cilostazol. n= 3 – 4 skin samples per mouse, n= 3 - 4 mice. Data represent the mean value ± SEM. *p< 0.05; **p < 0.01.

We next explored whether the induction of HF growth induces the transitory increase in the caliber of LV. To this end, to avoid severing LV and the induction of an inflammatory response, we did not conduct hair plucking experiments to synchronize HF, but rather stimulated the Telogen to Anagen transition by reducing the number of perifollicular macrophages at early Telogen with Clodronate liposomes (Fig 4C and D), as previously described (Castellana *et al.*, 2014). Under these conditions, LV surrounding the precocious Anagen HF displayed an increase in their caliber compared to controls (Fig 4C and D). We further analyzed the connection between HFSC cell proliferation and the expansion of LV caliber in K14Cre^+/T^, ∆Nβ-catenin^lox/lox^ mouse skin samples. This model is characterized by continuous HFSC proliferation and pilomatricoma formation (Jensen *et al.*, 2009; Lowry *et al.*, 2005). Immunofluorescence analyses for LYVE1 revealed a significant increase in LV caliber under conditions of continuous HF growth (Fig 4E and F). To interrogate if a sustained increased in vascular flow prompts the entry of Telogen HF into Anagen, we pharmacologically treated mice in early Telogen (P49) with Cilostazol (Kimura *et al.*, 2014); as previously documented (Choi *et al.*, 2018), increasing the vascular flow led to precocious HF growth and an increase of in the percentage of HF in anagen (Fig 4G). Overall, these results exposed the existence of networks of LV that dynamically reorganize increasing their caliber upon activation of HFSC.

### Transcriptome analysis of lymphatic endothelial cells at the onset of HFSC activation

Our prior results prompted us to analyze the existence of a differential gene expression in LV during the physiological HF Telogen to Anagen transition, when LV exhibit an expanded caliber (Fig 4A and B). We focused on the physiological Tm (P55), and Tl (P70) phases of the second HF cycle as HF progress to Anagen. To this end, we FACS-isolated eYFP+ dermal LV from the backskin of Prox1CreERT2^+/T^; Rosa26-LSL-eYFP^KI/KI^ mice, where 60% of the represent lymphatic endothelial cells (Bianchi *et al.*, 2015). We avoided including the dermal subcutaneous collector LV, through mechanical removal before tissue digestion. In order to comprehensively define changes in the molecular traits of isolated cells, we conducted RNA sequencing (RNA seq) analyses. The transcriptome profiles revealed the expression of genes differentially regulated between the analyzed populations. We first determined the existence of differentially expressed pathways in Tl compared to Tm phases, establishing LV gene expression rankings using gene set enrichment analyses (GSEA) compared with public annotations from Reactome, Biocarta, Kyoto Encyclopedia of Genes and Genomes (KEGG) and Gene Ontology (GO) databases. The majority of the differentially regulated genes (FDR q-values < 0.25) were related to cell adhesion, cytoskeleton and lymphatic organization/axon guidance (Fig 5A). Further analyses revealed that 310 genes were upregulated and 562 downregulated with a 2.8-fold change (log2 fold change > 1.5 or < −1.5) at Tl (P70) compared to Tm (P55). Enrichr analyses (Chen *et al.*, 2013; Kuleshov *et al.*, 2016) were then used to classify the major represented categories, which exposed the differential expression of genes related to LV remodeling (Fig 5B & 5C), in agreement with the transitory morphological changes observed in LV at the onset of HFSC activation. The genes encoded proteins involved in ECM, cytoskeleton and adhesion processes and vesicular trafficking, including the upregulation of ECM proteins (e.g. Tenascin X, TNXB; heparan sulfate proteoglycan 2, HSPG2; Collagen V type 3 chain, COL5A3), actin and microtubule organization (e.g. Rho guanine nucleotide exchange factor 1, ARHGEF1; Plectin, PLEC; Dynactin, DCTN1; Kinesin family members, KIF22 & KIF26A), and adhesion proteins (e.g. Platelet endothelial cell adhesion molecule, PCAM1/CD31; Plakoglobin, JUP; Integrin Subunits Alpha 5 and 6, ITGA5 & ITGA6; Basal Cell Adhesion Molecule, BCAM). Also, genes involved in lymphatic remodeling and axon guidance were found upregulated (e.g. Endoglin, ENG; Semaphorins; SEMA3F & SEMA4D; EMILIN1; Plexin D1, PLXND1; Polycystin-1, PKD1; Netrin Receptor 5A, UNC5A) as well as genes involved in intracellular signaling processes such as upregulation in transforming growth factor B1 (TGFB1) or downregulation of chemokines and cytokines (e.g. CXCL1, 2, 7, 19, CCL5, 8, 10, 11, 22, IL2,6). These genes have been involved in the regulation of lymphatic remodeling, lumen organization and permeability, and taken together these results support the existence of a transitory dynamic reorganization of LV at the onset of HFSC activation.

**Figure 5.**
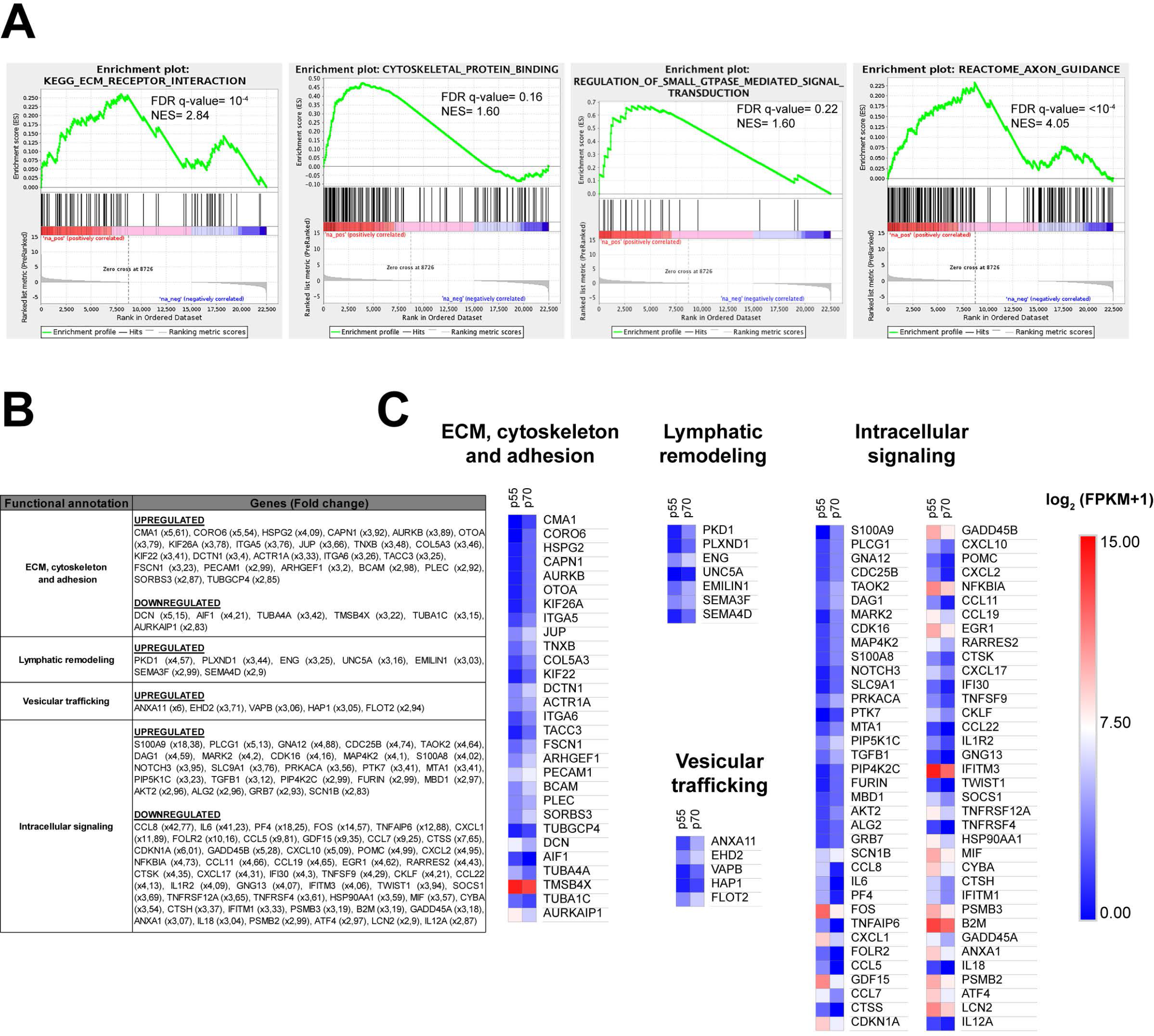
Transcriptome analysis of LV in late Telogen compared to mid Telogen. A. Gene set enrichment plots (GSEA) of LV using public annotations showing the enrichment of selected gene signatures involved in different pathways in late Telogen (P69, Tl) compared with mid Telogen (P55, Tm). NES: normalized enrichment score. The false discovery rate (FDR; q-value) is indicated. B. Table of differentially expressed genes (DEGs), in LV during late Telogen compared to mid Telogen, relative to biological processes, analyzed using the Enrichr platform (Chen *et al.*, 2013; Kuleshov *et al.*, 2016). Upregulated (red), downregulated (blue). Data shown represent the fold change of the genes. C. Heat map diagram of DEGs in LV during late Telogen compared to mid Telogen, relative to biological processes, analyzed using the Enrichr platform (Chen *et al.*, 2013; Kuleshov *et al.*, 2016). Data shown represent the log2 (FPKM+1).

### Depletion of LV blocks the pharmacological induction of HF growth

To get insight into the functional association between LV and the HF cycle, we pharmacologically stimulated HF growth, followed by loss of function approaches using the mouse genetic model Prox1CreERT2^+/T^;Rosa26-LSL-iDTR^KI/KI^ mice to conditionally ablate LV. Prox1CreERT2^+/T^;Rosa26-LSL-iDTR^KI/KI^ and controls were first treated at early telogen (P49) with Cyclosporine A (CSA), a potent hair growth stimulator (Gafter-Gvili *et al.*, 2003; Horsley *et al.*, 2008; Paus *et al.*, 1989); which activates quiescent HFSC through the inhibition of calcineurin and the transcription factor of activated T cells c1 (NFATc1) (Gafter-Gvili *et al.*, 2003; Horsley *et al.*, 2008). Consistent with those prior findings, the precocious growth of HF in CSA treated mice was fully distinctive at the end of the experiment compared to controls, as observed by histological approaches (Fig 6A and B). Precocious HF exhibited a significant increase in proliferation, as evidenced by immunohistological analyses of the proliferation marker Ki67 (Fig 6C and D). Moreover, in agreement with our prior findings (Fig 4) the induction of precocious HF growth promoted a significant increase in LV caliber (Fig 6E and F).

**Figure 6.**
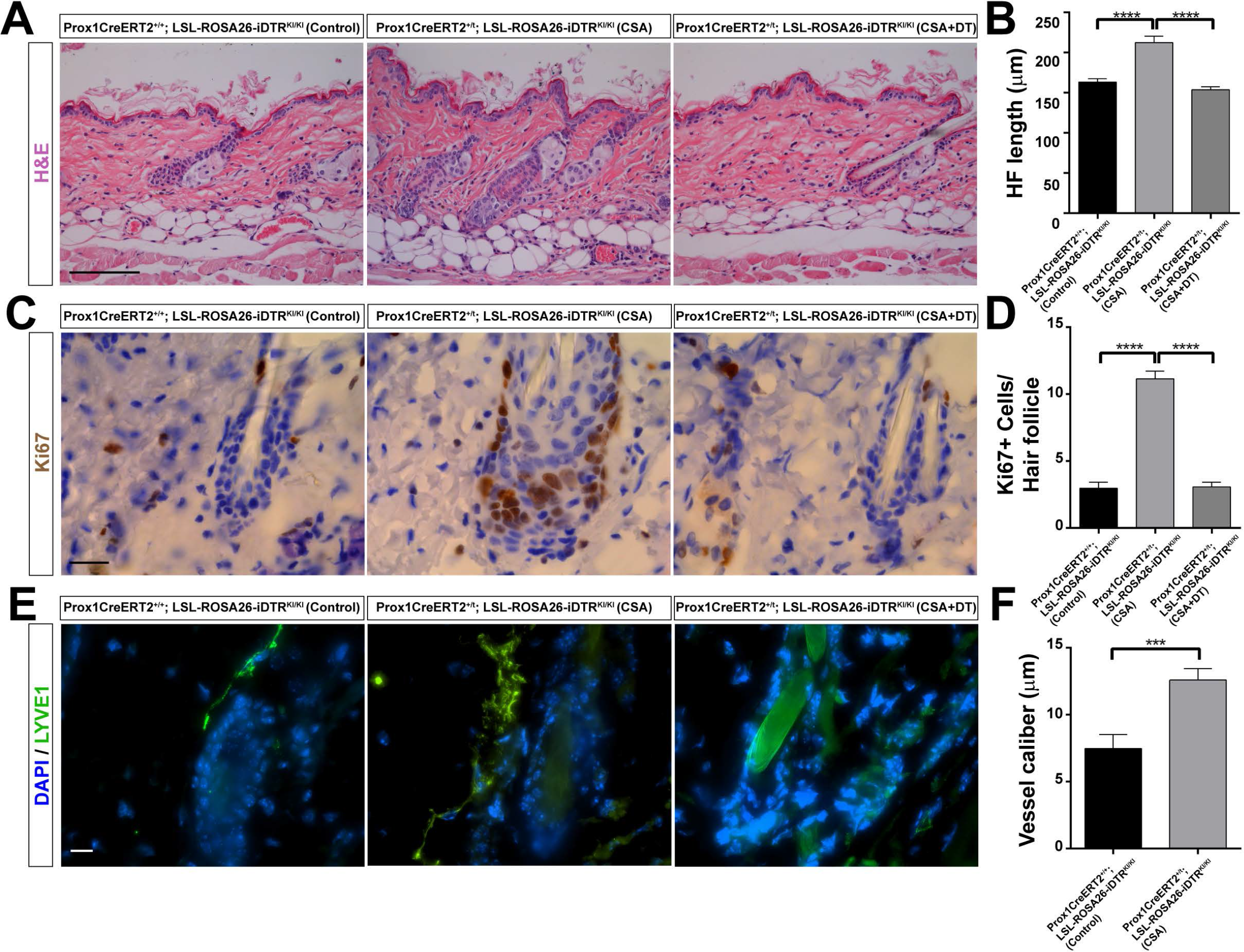
Depletion of LV blocks the pharmacological induction of HF growth. A. H&E staining of adult backskin sections from Prox1CreERT2^+/+^; LSL-ROSA26-iDTR^KI/KI^ injected with vehicles (Control), Prox1CreERT2^+/T^; LSL-ROSA26-iDTR^KI/KI^ mice treated with CSA and vehicle (CSA); and Prox1CreERT2^+/T^; LSL-ROSA26-iDTR^KI/KI^ treated with tamoxifen, CSA and intradermal Diphtheria Toxin (CSA+DT). n= 3 – 4 skin samples per mouse, n= 3 - 4 mice. Bar, 200 μm. B. Histogram of the HF length in the skin of Prox1CreERT2^+/+^; LSL-ROSA26-iDTR^KI/KI^ injected with vehicles (Control), Prox1CreERT2^+/T^; LSL-ROSA26-iDTR^KI/KI^ mice treated with CSA and vehicle (CSA); and Prox1CreERT2^+/T^; LSL-ROSA26-iDTR^KI/KI^ treated with tamoxifen, CSA and intradermal Diphtheria Toxin (CSA+DT). n= 3 – 4 skin samples per mouse, n= 3 - 4 mice. Data represent the mean value ± SEM. *p< 0.05; **p < 0.01; ***p < 0.001. C. Ki67 immunostaining of adult backskin sections from Prox1CreERT2^+/+^; LSL-ROSA26-iDTR^KI/KI^ mice injected with vehicles (Control), Prox1CreERT2^+/T^; LSL-ROSA26-iDTR^KI/KI^ mice treated with CSA and vehicle; and Prox1CreERT2^+/T^; LSL-ROSA26-iDTR^KI/KI^ treated with tamoxifen, CSA and intradermal Diphtheria Toxin. n= 3 – 4 skin samples per mouse, n= 3 – 4 mice. Bar, 20 μm. D. Histogram of the number of Ki67 positive cells per HF in skin sections from Prox1CreERT2^+/+^; LSL-ROSA26-iDTR^KI/KI^ injected with vehicles (Control), Prox1CreERT2 ^+/T^; LSL-ROSA26-iDTR^KI/KI^ mice treated with CSA and vehicle (CSA); and Prox1CreERT2^+/T^; LSL-ROSA26-iDTR^KI/KI^ treated with tamoxifen, CSA and intradermal Diphtheria Toxin (CSA+DT) n= 3 – 4 skin samples per mouse, n= 3 - 4 mice. Data represent the mean value ± SEM. *p< 0.05; **p < 0.01; ***p < 0.001. E. Adult backskin sections from Prox1CreERT2^+/+^; LSL-ROSA26-iDTR^KI/KI^ injected with vehicles (Control), Prox1CreERT2^+/T^; LSL-ROSA26-iDTR^KI/KI^ mice treated with CSA and vehicle (CSA); and Prox1CreERT2^+/T^; LSL-ROSA26-iDTR^KI/KI^ treated with tamoxifen, CSA and intradermal Diphtheria Toxin (CSA+DT) immunostained for LYVE1 (green), and counterstained with DAPI (blue). Bar, 10 μm. n= 3 – 4 skin samples per mouse, n= 3 - 4 mice. F. Histogram of the LV caliber in skin sections from Prox1CreERT2^+/+^; LSL-ROSA26-iDTR^KI/KI^ injected with vehicles (Control), Prox1CreERT2^+/T^; LSL-ROSA26-iDTR^KI/KI^ mice treated with CSA and vehicle (CSA); and Prox1CreERT2^+/T^; LSL-ROSA26-iDTR^KI/KI^ treated with tamoxifen, CSA and intradermal Diphtheria Toxin (CSA+DT) n= 3 – 4 skin samples per mouse, n= 3 - 4 mice. Data represent the mean value ± SEM. *p< 0.05; **p < 0.01; ***p < 0.001.

CSA treated Prox1CreERT2-iDTR mice were further treated with intradermal injections of diphtheria toxin for 3 d before the end of the CSA treatment. This treatment led to a significant ablation of LV, as confirmed by reductions in lymphatic drainage, supported by the impairment of Evans Blue dye clearance in skin (Fig EV3A), and LYVE-1 immunofluorescence analyses (Fig EV3B). Strikingly, the loss of LV precluded the precocious activation of HF growth induced by CSA, as evidenced by histological features (Fig 6A and B) and a significant reduction of Ki67 proliferating cells (Fig 6C and D), in comparison to CSA treated controls. As an additional control, we also included Prox1CreERT2^+/T^; Rosa26-LSL-iDTR^KI/KI^ mice at early telogen that were not treated with CSA, and no changes in HF organization of growth were observed upon LV ablation with diphtheria toxin (Fig EV4). These results exposed a functional connection between LV upon induction of the early onset of HF regeneration.

## Discussion

Mammalian HF undergo a lifetime cyclic regeneration, and several discoveries have increased our understanding on the regulation of this significant model of organ regeneration, both at individual HF level, through coordinated interactions with other cells at the HFSC niche or molecular signaling across adjacent HF across the tissue (Gonzales & Fuchs, 2017; Guasch, 2017; Widelitz & Chuong, 2016). Here, we now show that within a single HF, HFSC associate with LV capillaries, starting from developmental HF stages. We further uncovered the existence of a dynamic patterned association of LV with adjacent HF across the skin, and defined that the loss of LV abrogates the pharmacological induction of HF regeneration (Fig EV5). These findings position LV as new elements of the HFSC niche, and open new interesting questions about the potential roles of LV in regulating individual and adjacent HF at different stages of the physiological HF cycle.

We first discovered that the interaction of LV with individual HF is polarized, and occurs along the anterior permanent HF region (Fig 1). Of note, we did not observe a preferential association of LV with particular HF types, as it has been observed for mechanosensory neurons (Li *et al.*, 2011). It remains to be investigated whether the polarized LV – HF association is dependent on the existence of differentially expressed molecules at the anterior/posterior sides of the HF, thereby creating a distinct connection similar to the one documented between HF and the apm (Fujiwara *et al.*, 2011). The nature of the lymphatic endothelial cells associated with HF is also intriguing, since they are plastic and heterogeneous among organs (Jang *et al.*, 2013; Martinez-Corral *et al.*, 2015; Petrova & Koh, 2018). Thus, a defined population may exert specific functions at HF, compared to lymphatic endothelial cells distributing at other anatomical locations within the skin.

It is generally well-acknowledged that HFSC create a special niche microenvironment, where the interactions with other cells within the niche are important for HFSC organization and function (Gonzales & Fuchs, 2017; Guasch, 2017). In this regard, we observed that a functional HFSC niche is required to sustain the polarized LV-HF association in the backskin, through the expression of Wnt ligands. In the absence of HFSC-derived Wnts, LV dissociate from HF allocating parallel to the epidermis (Figure 1G and H), in a similar disposition to the one observed in the ear skin (Fig EV1A), supporting the existence of unique LV regional microenvironments. Interestingly, the lymphatic capillaries that associate to individual HF further branch at the level of HFSC, converging into a single LV for every three HF units, which in turn interconnect with other HF triads across the skin (Fig 3). This patterned LV organization resembles the one documented for dermal blood vessels distributed at the dermal plexus (Sada *et al.*, 2016). At this location, blood vessels set different regional areas and contribute to the maintenance of two independent epidermal stem cell populations. However, respecting HFSC, blood vessels are distributed in a distinct manner, forming a venule annulus (Xiao *et al.*, 2013). It remains to be explored whether LV exert a defined regulatory role on a particular SC population, since distinct populations of SC exhibiting different potential localize along the HF permanent region and at the HF bulge (Guasch, 2017).

At tissue level, we observed that LV dynamically flow through neighboring HF across the skin. Interestingly, the LV caliber transiently expanded at the onset of HFSC activation (Fig 4), suggesting an increased tissue drainage capacity. Studies combining live imaging approaches and biophysical analyses of LV will be instrumental for defining spatiotemporal changes in LV dynamics at different stages of the HF cycle. The dynamic flow of LV through HF likely facilitates the distribution of molecules and immune cells across the skin. Although it remains unclear whether changes in the diffusion of soluble or cellular components through LV is linked to HF cycle regulation, our results now add LV as new roads connecting HF across the skin and pave the way for further investigation. In this regard, it is well acknowledged that the coordinated propagation of stimulatory/inhibitory signals across the skin regulates the HF cycle (Widelitz & Chuong, 2016). Interestingly, measurements of the decline levels of diffusible molecules, including growth factors, morphogens, cytokines, predicted that their range of action is restricted to no more than 100 μm (Teleman & Cohen, 2000; Weber *et al.*, 2013). However, in the backskin it has been observed that the effect of diffusible molecules is exerted at more than 1 mm further from their initial expression site (Chen *et al.*, 2015). Thus, LV could potentially partake in this event aiding the diffusion of molecules at longer distances. Also, LV functional roles in draining tissue fluids, could further modulate the concentration level of molecular inhibitors or activators at specific phases of the HF cycle.

LV may also regulate immune cell trafficking to HF areas, although immune privilege areas are normally devoid of LV (Paus *et al.*, 2003). Indeed, it has been recently reported that Foxp3 + T regulatory cells, which traffic through LV (Hunter *et al.*, 2016), localize to HF and regulate HFSC proliferation (Ali *et al.*, 2017). Perifollicular macrophages have also been shown to contribute to the activation of HFSC through the expression of Wnt ligands (Castellana *et al.*, 2014), and myeloid Wnt ligands regulate the development of LV(Muley *et al.*, 2017). Moreover, macrophages regulate LV caliber (Gordon *et al.*, 2010), through the expression of Wnt ligands (Muley *et al.*, 2017). Interestingly, our observations on the transient increment of the LV caliber at the onset of HFSC activation, also coincide with the documented expression of macrophage-derived Wnt ligands and the activation of HFSC (Castellana *et al.*, 2014). Thus, changes in the LV caliber could exert regulatory functions through the spatiotemporal diffusion of morphogens and the influx and outflow of immune cells to HF at specific phases of the HF cycle.

Our results exposed a marked increase of the LV caliber during the physiological late telogen to anagen transition (Fig 4). The induction of HFSC activation by modulating macrophage levels or upon pharmacological induction with CSA (Fig 4 and Fig 6) exposed a functional connection between HFSC and LV caliber expansion upon HFSC activation. These changes were sustained under conditions of continuous HF growth as observed in the skin of K14Cre^+/T^, ∆Nβ-catenin^lox/lox^ mice (Fig 4). During the physiological telogen to anagen transition these LV changes were associated with a more irregular and fenestrated lumen morphology, in agreement with the distinct expression of genes encoding proteins mainly involved in lymphatic cytoskeletal remodeling and matrix cell adhesion (Fig 5A and B), suggesting the occurrence of mechanotransduction processes, changes in interstitial pressure and modulation of tissue drainage (Planas-Paz & Lammert, 2013; Sabine *et al.*, 2016). Whether these modifications could also lead to changes in the HFSC niche stiffness, which are known to regulate HFSC behavior (Lane *et al.*, 2014), needs further investigation. The physiological roles of the identified LV signatures, as well as their potential roles in the activation/inhibition of HF regeneration as direct/indirect effectors remains to be explored. To gain insight into the functional connection of LV in the regulation of HF behavior, we conditionally depleted LV upon the pharmacological induction of HFSC activation. These transient analyses exposed a functional connection between LV and HFSC regeneration upon induction of the early onset of HF growth (Fig 6).

In future studies, it will be important to analyze the contributions of LV along the different stages of the physiological HF cycle as well as in other specific HF stages. The study of the contribution of LV at defined HF stages is particularly relevant, since the temporal diffusion of activators or inhibitors must likely lead to different responses in HF cycling in a spatiotemporal manner. Taken together, our results position LV as novel components of the HFSC niche coordinating HF connections at tissue-level and provide insight into their functional association to the HF cycle.

## Experimental Procedures

### Mice and treatments

All mouse experiments were approved and performed according to Institutional and ethical regulations of the CNIO and the Institute of Health Carlos III.

Backskins from C57Bl/6 mice (n=3-5) from different embryonic (E15.5, E16.6 and E17.5) and postnatal (P) days (P5, P12, P16, P23, P35, P45, P49, P55, P69 and P85) were collected to analyze different phases of the HF cycle.

The Prox1-CreERT2 mouse model (Tg(Prox1-cre/ERT2)#aTmak, a kind gift from Dr. Taija Mäkinen, Uppsala University)(Bazigou *et al.*, 2011) was crossed under the background of the ROSA26-LSL-eYFP reporter mice. The eYFP expression was induced by intraperitoneal injections of 2 mg Tamoxifen (T5648, Sigma-Aldrich) in sunflower oil for 4 d.

Skin resident macrophages were reduced by treating P49 mice with intradermal injections of 1 mg of Clodronate encapsulated liposomes or empty liposomes as controls (Encapsula Nanosciences), as described previously (Castellana *et al.*, 2014).

CSA treatments were carried out by treating P49 mice with intraperitoneal injections of 100mg/kg CSA (30024, Sigma-Aldrich) or vehicle control for 10 d before sacrifice.

Cilostazol treatments were carried out by treating P49 mice with intradermal injections of 100 µg Cilostazol or vehicle control every two days for 9 d before sacrifice.

The Prox1-CreERT2; ROSA26-LSL-iDTR (Gt(ROSA)26Sortm1(HBEGF)Awai/J, Jackson) mice were used to ablate LV upon induction of HF growth. To this end, mice were first treated with the intraperitoneal administration of 100 mg/kg Cyclosporin A (30024, Sigma-Aldrich) during 10 d starting at P49. At P52, mice were injected intraperitoneally with 2 mg Tamoxifen in sunflower oil for 4 d to induce the expression of the diphtheria toxin receptor, followed by the intradermal administration of 1ng/g diphtheria toxin (322326, Calbiochem) for 3 d starting at P56 before sacrifice at P58. To assess LV ablation, mice were intradermally injected with 10 μl of 1% Evans blue dye (E2129, Sigma-Aldrich). Skin samples were collected 16 h later and incubated in formamide (F9037, Sigma-Aldrich) at 55º for 24 h to extract the dye from the tissue. The dye levels were quantified by measuring the absorbance of the dye at 610 nm.

K15-CrePR1 mice [(Krt1-15-cre/PGR)22Cot/J](Morris *et al.*, 2004), and Wlstm^1.1Lan/J^ mice (Carpenter *et al.*, 2010) were acquired from Jackson Labs. K15-CrPR1^+/T^/; Wls^lox/lox^ 7d old mice were then treated with Mifepristone (#M8046, Sigma-Aldrich) administered in the drinking water at 429 ng/mL (Babij *et al.*, 2003) to the mother females and continued in weaned mice until the end of the experiment (12 weeks of treatment). K14∆Nβ-cateninER-transgenic mouse backskin samples were kindly provided by Dr. Kim Jensen (Biotech Research and Innovation Centre - BRIC, University of Copenhagen)(Jensen *et al.*, 2009).

### Immunofluorescence

Backskin samples were embedded in paraffin blocks and Optimal Cutting Temperature (OCT) compound. Paraffin sections were rehydrated and treated for antigen retrieval using 10 mM Sodium Citrate, 0.05% Tween 20, pH 6.0 at 100°C for 10 min. Slides were incubated in blocking buffer: 0.3% Triton X100 – PBS (PBST), 5% FBS (F7524, Sigma-Aldrich), 1% gelatin from cold water fish skin (G7765, Sigma-Aldrich) and 1% BSA (10 735 078 001, Roche). Primary antibodies (Table S1) were diluted in blocking buffer and incubated overnight at 4°C and washed. The slides were then incubated for 1 h with the corresponding secondary antibodies (Table S2) and counterstained with DAPI. For OCT sections, samples were fixed in 4% PFA for 10 min and the antibody stainings were performed as described for paraffin embedded sections.

For whole mounts and tissue clearing, embryo and postnatal backskins were fixed overnight in 4% PFA (15710, Electron Microscopy Sciences) at 4°C. The backskins were cut into small pieces, washed 10 times in PBST for 30 min each, and incubated with primary antibodies diluted in Blocking Buffer (0.3% PBST, 5% FBS and 20% DMSO) at room temperature during 5 d. Backskins were washed in PBST as previously described, followed by their incubation with secondary antibodies and counterstained with DAPI for 3 d at room temperature. Tissues were dehydrated in increasing concentrations of methanol (25%, 50%, and 75% methanol for 5 min) and placed in 100% methanol 3 times 20 min each. Next, backskins were cleared overnight in 1:2 benzyl alcohol: benzyl benzoate (BABB) (B6630 and 402834, Sigma-Aldrich).

Images were acquired in a TCS-SP5 (AOBS) confocal (Leica microsystems) microscope. High-resolution images were captured using a 63X HCX PL APO 1.3 Glycerol immersion objective.

### Intravital confocal microscopy

Mice were anesthetized with 1.5% isoflurane and placed into a custom-made adapter with a stable temperature during the whole duration of the experiments. Intravital experiments were carried out in a TCS-SP5 (AOBS) confocal (Leica microsystems). The image acquisition was conducted during 8 h every 15 min, using a 20x HCXPL APO 0.7 N.A. dry objective.

### FACS and RNA extraction

Backskins of Prox1-CreERT2; ROSA26-LSL-eYFP mice were collected the day after the last intraperitoneal injection of 2 mg Tamoxifen. Next, the subcutaneous fat was removed with a scalpel, and the skin samples were cut into small pieces and incubated in 0.33 mg/ml of Liberase TM Research Grade (05401119001, Roche) at 37 °C for 30 min. After incubation, skin samples were transferred to a MACS gentle dissociator C tube (130-093-237, Miltenyi biotec), and filtered through 40 and 70 μM cell strainers (352350 and 352340, Falcon). Red blood cells were eliminated with lysing buffer (555899, BD Biosciences). DAPI was added as a viability dye, and FACS-isolated eYFP positive cells (FACS ARIA IIu sorter, Becton Dickinson) were collected in Lysis Buffer (400753, Absolutely RNA Nanoprep kit, Agilent Technologies).

For the analysis of the expression of Wls in HFSC, HFSC were FACS isolated and stained as previously described using the markers CD34 and α6 integrin (Castellana *et al.*, 2014). RNA extraction was performed following manufacturer’s instructions.

### RNA isolation and Real Time-PCR (RT-PCR)

1 μg of total RNA isolated from skin using TRIZOL (15596026, Invitrogen) was used for cDNA synthesis using the Ready-to-Go You-Prime It First-Strand beads and random primers (GE Healthcare). RT-PCR reactions were conducted using the GoTaq qPCR Master Mix (A6001, Promega) and a MasterCycler Ep-Realplex thermal cycler (Eppendorf).

The expression levels were normalized to Actin. The complete list of primers used is presented in Table S3.

### RNA sequencing

Variable amounts of total RNA samples, between 50 and 2,000 pg, were processed with the SMART-Seq v4 Ultra Low Input RNA Kit (Clontech) by following manufacturer instructions. Resulting cDNA was sheared on an S220 Focused-ultrasonicator (Covaris) and subsequently processed with the "NEBNext Ultra II DNA Library Prep Kit for Illumina" (NEB #E7645). Briefly, oligo(dT)-primed reverse transcription was performed in presence of a template switching oligonucleotide, double stranded cDNA was produced by limited-cycle PCR and submitted to acoustic shearing. Fragments were processed through subsequent enzymatic treatments of end-repair, dA-tailing, and ligation to Illumina adapters. Adapter-ligated libraries were completed by 6 cycles of PCR. Reads were sequenced in single-end mode (51 bp) on an Illumina HiSeq2500 by following manufacturer’s protocols.

### Gene expression analysis by RNA-seq and Gene set enrichment analysis (GSEA)

The differential expression of genes was assessed using the Nextpresso pipeline (http://bioinfo.cnio.es/nextpresso/). The sequencing quality was analyzed with FastQC (http://www.bioinformatics.babraham.ac.uk/projects/fastqc/); reads were aligned to the mouse genome (GRCm38/mm10) using TopHat-2.0.10 (Trapnell *et al.*, 2012), Bowtie 1.0.0 (Langmead *et al.*, 2009) and Samtools 0.1.19.0 (Li *et al.*, 2009). The assembly of transcripts, abundance estimation and differential expression were calculated with Cufflinks 2.2.1 (Trapnell *et al.*, 2012), using the mouse GRCm38/mm10 transcript annotations from https://ccb.jhu.edu/software/tophat/igenomes.shtml. We then used the RNA-seq gene list, where genes were ranked according to their statistical expression difference to perform a GSEAPreranked analysis (Subramanian *et al.*, 2005), using pathway annotations from Reactome, Biocarta, NCI (http://www.ndexbio.org/#/user/301a91c6-a37b-11e4-bda0-000c29202374) and KEGG public databases.

All basic and advanced fields were set to default and we considered only the gene sets with False Discovery Rate (FDR) q-values < 0.25 that were significantly enriched.

For enrichment analysis, genes with a variation of log2 fold change > 1.5 or < −1.5 were selected and introduced in the Enrichr tool (Chen et al., 2013; Kuleshov et al., 2016). The genes that belonged to the categories of interest were chosen for the heatmaps. Heatmaps representing the log2(FPKM+1) levels were done using the software Morpheus (https://software.broadinstitute.org/morpheus).

### Statistical Analyses

Image analyses were performed using Image J and Imaris software (Bitplane Scientific Software, Zurich). For statistical analysis of quantitative data, the data normality was evaluated and data that presented a Gaussian distribution was analyzed using two-tailed Student’s t-test. Statistical analyses were done using GraphPad Software (La Jolla, Ca). All data is representative of at least two independent experiments performed in triplicates.

## Author Contributions

D.P-J. and S.F. experimental design and analyzes. D.M. confocal and intravital microscope studies, C.F and O.G. RNA seq methodological analyses, D.C. samples and experimental analyses, and R.L. reagents, hypotheses discussion and input. M.P-M study conception, supervision and funding. D.P-J., and M.P-M. manuscript writing with input from all authors.

## Acknowledgments

We thank Dr. Kari Alitalo (University of Helsinki) for advice, Dr. Taija Mäkinen (Uppsala University) for providing the Prox1ERT2 mice, Dr. Kim Jensen (BRIC, University of Copenhagen for providing K14Cre^+/T^, ∆Nβ-catenin^lox/lox^ mouse skin samples, and Susan Morton for the Lhx2/9 antibody (T. Jessel lab, Columbia University). We thank the core facilities of the CNIO and the University of Copenhagen for technical assistance. This work was supported by grants from the Spanish Ministry of Economy and Competitiveness/European Regional Development Fund (ERDF), European Union (BFU2015-71376-R to M.P-M), the Worldwide Cancer Research UK Foundation (15-1219 to M P-M) and the Novo Nordisk Foundation (NNF17OC0028028). The authors declare no conflict of interest.

## Figures - Expanded View

**Figure EV1.**
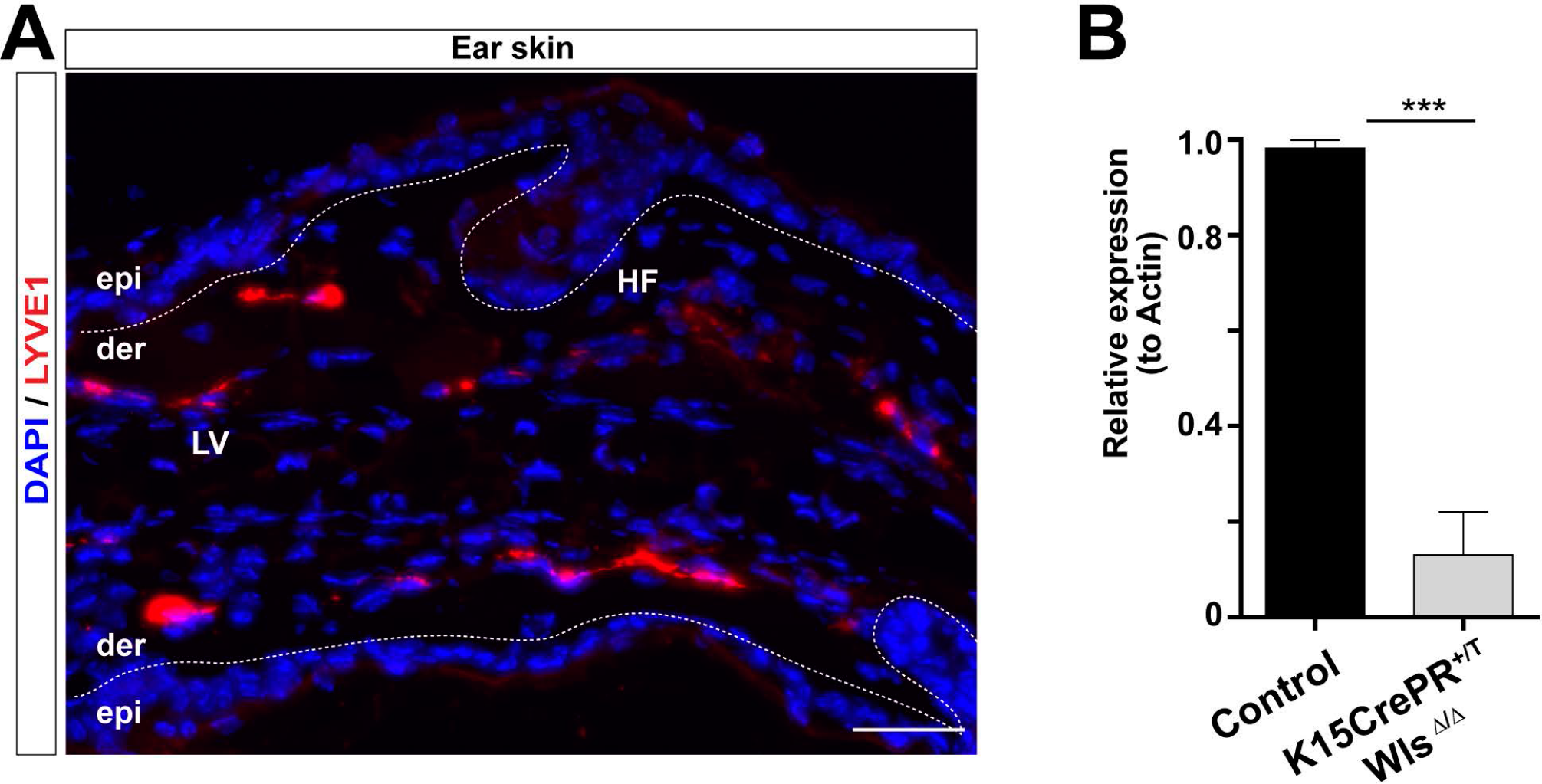
Distribution of LV in the ear skin and validation of the reductions of Wls expression in the K15CrePR^+/T^; Wls^Δ/Δ^ mouse model. A. Adult ear skin sections immunostained for LYVE1 (red), and counterstained with DAPI (blue). n= 3 – 4 skin samples per mouse, n= 3 - 4 mice. Bar, 50 μm. LV, lymphatic vessels; HF, Hair follicles; epi, epidermis; der, dermis. B. Histogram of the RT-qPCR analyses of the relative expression of Wls in the HFSC isolated from the K15CrePR^+/T^; Wls^Δ/Δ^ mouse model and controls. Data represent the mean value ± SEM. ***p < 0.001.

**Figure EV2.**
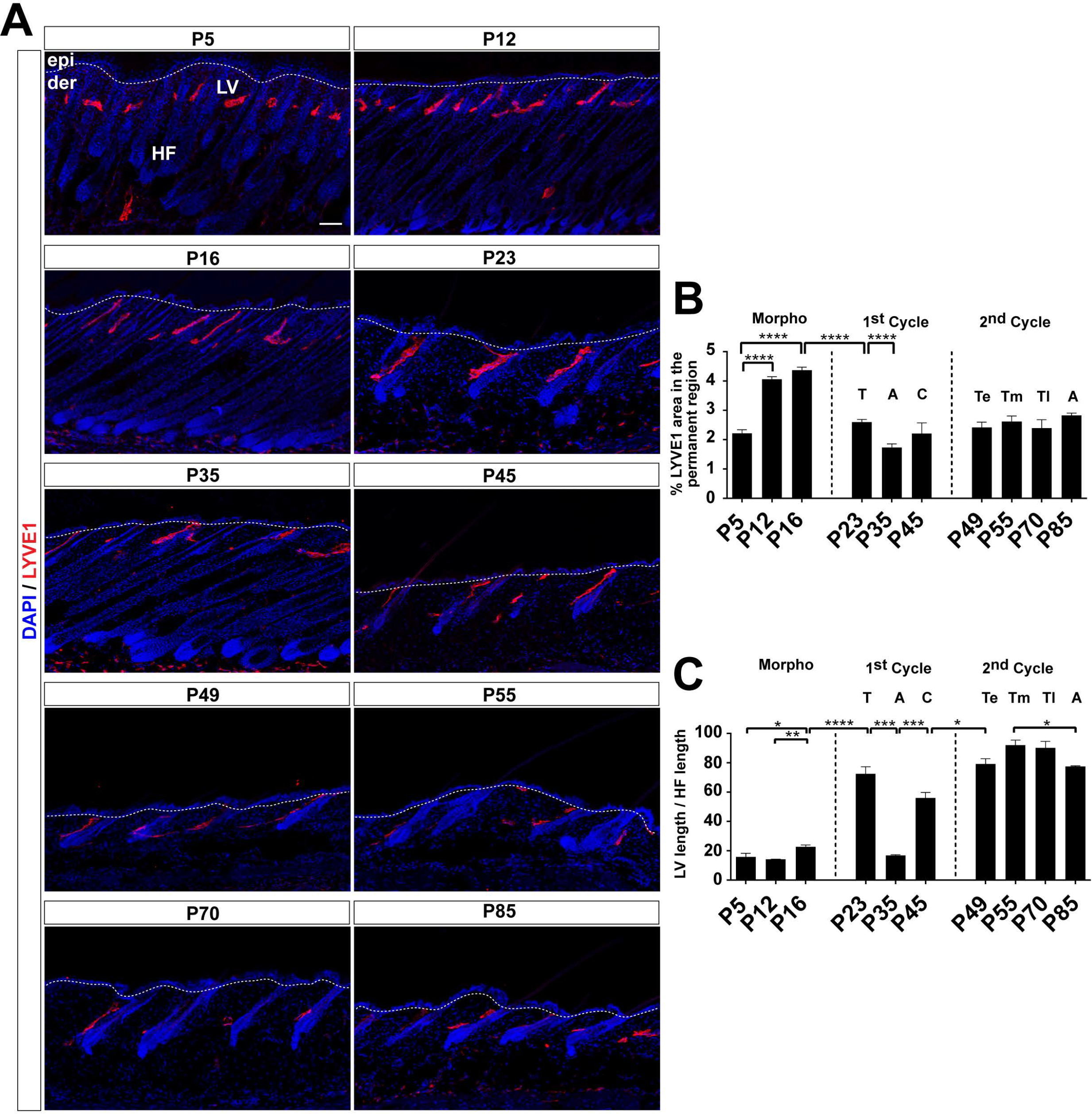
LV association with HF during the postnatal HF cycle. A. Adult backskin sections from different postnatal (P) days immunostained for LYVE1 (red) and counterstained with DAPI (blue). n= 3 – 4 skin samples per mouse, n= 3 - 4 mice. Bar, 50 μm. epi, epidermis; der, dermis; LV, lymphatic vessels; HF, hair follicle. B. Histogram of the percentage of LYVE1 positive area in the HF permanent region at different postnatal days. n= 3 – 4 skin samples per mouse, n= 3 - 4 mice. A, Anagen; C, Catagen; T, Telogen; Te, early Telogen; Tm, mid Telogen; Tl, late Telogen. C. Histogram of the percentage of LV length relative to the HF length in the backskin at different postnatal days. n= 3 – 4 skin samples per mouse, n= 3 - 4 mice. A, Anagen; C, Catagen; T, Telogen; Te, early Telogen; Tm, mid Telogen; Tl, late Telogen. The data shown in all histograms represent the mean value ± SEM. *p< 0.05; **p < 0.01; ***p < 0.001.

**Figure EV3.**
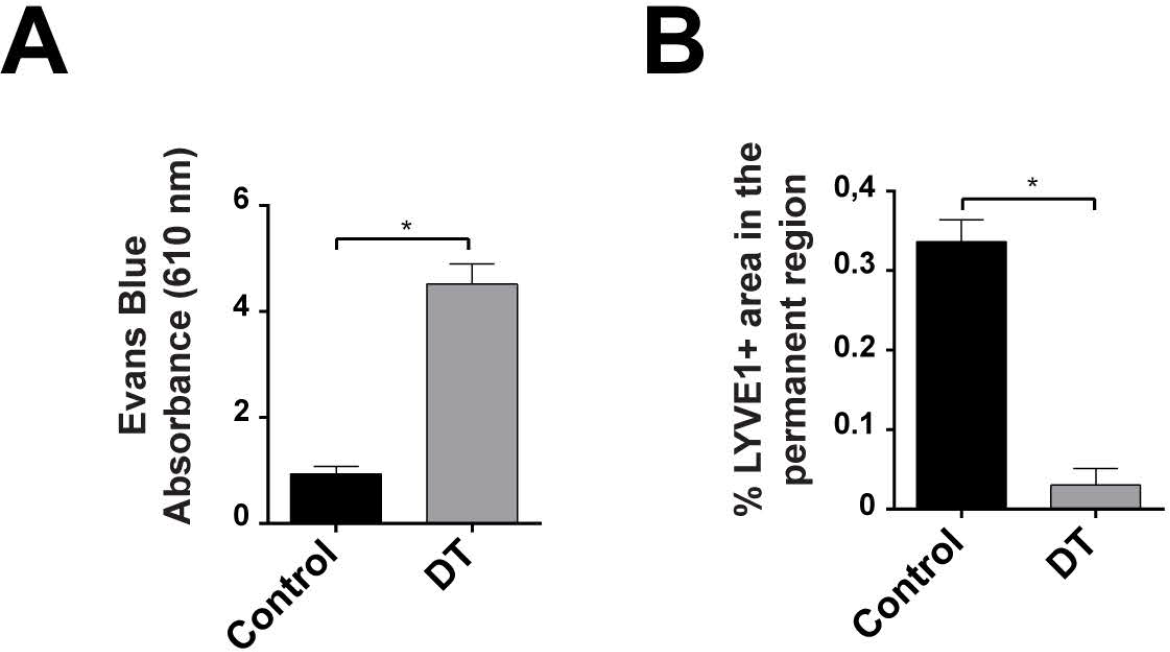
Characterization of the efficiency of LV depletion in the backskin of Prox1CreERT2 ^+/T^; LSL-ROSA26-iDTR^lox/lox^ mice upon treatment with diphteria toxin. A. Histogram of the concentration of Evans Blue (absorbance 610 nm) in the skin of Prox1CreERT2^+/+^; LSL-ROSA26-iDTR^lox/lox^ mice (Control) and Prox1CreERT2 ^+/T^; LSL-ROSA26-iDTR^lox/lox^ mice treated intradermally with diphtheria toxin. n= 3 – 4 skin samples per mouse, n= 3 - 4 mice. Data represent the mean value ± SEM. *p< 0.05; **p < 0.01; ***p < 0.001. B. Histogram of the percentage of LYVE1 positive area in the HF permanent region in skin sections from Prox1CreERT2^+/+^; LSL-ROSA26-iDTR^lox/lox^ mice (Control) and Prox1CreERT2 ^+/T^; LSL-ROSA26-iDTR^lox/lox^ mice treated intradermally with diphtheria toxin. n= 3 – 4 skin samples per mouse, n= 3 - 4 mice. Data represent the mean value ± SEM. *p< 0.05; **p < 0.01; ***p < 0.001.

**Figure EV4.**
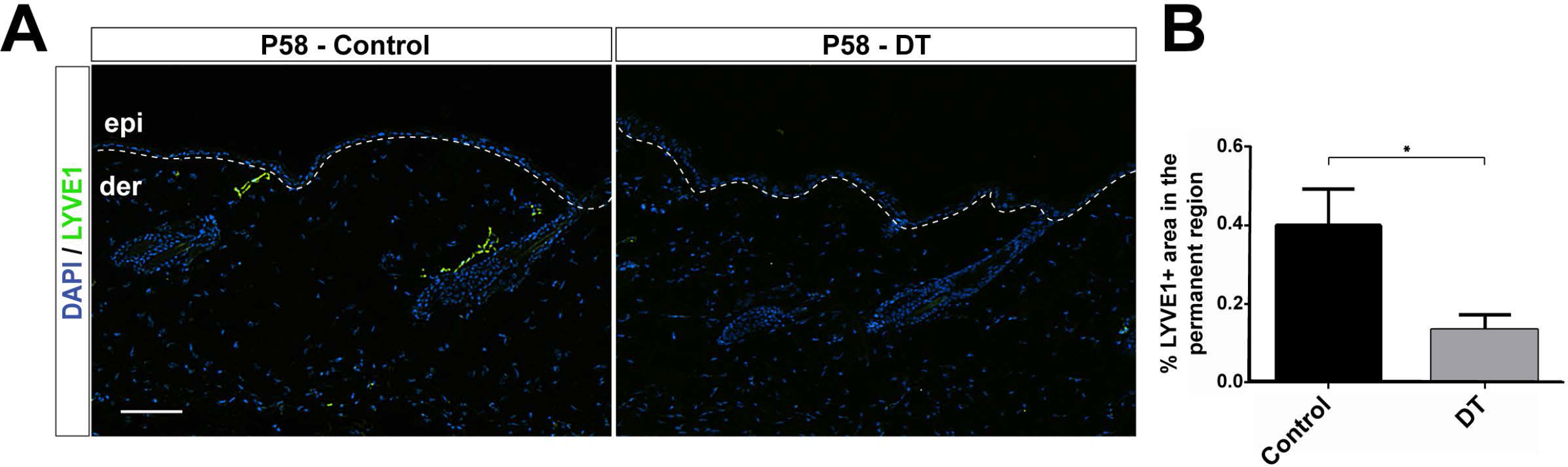
Depletion of LV during Telogen does not induce changes in HF organization or growth. A. Backskin sections from Prox1CreERT2^+/+^; LSL-ROSA26-iDTR^lox/lox^ injected with vehicle (Control), Prox1CreERT2^+/T^; LSL-ROSA26-iDTR^lox/lox^ mice treated with intradermal diphtheria toxin immunostained for LYVE1 (green), and counterstained with DAPI (blue). Bar, 10 μm. n= 3 – 4 skin samples per mouse, n= 3 - 4 mice. B. Histogram of the HF length in the skin of Prox1CreERT2^+/+^; LSL-ROSA26-iDTR^lox/lox^ injected with vehicle (Control), and Prox1CreERT2^+/T^; LSL-ROSA26-iDTR^lox/lox^ treated with intradermal diphtheria toxin. n= 3 – 4 skin samples per mouse, n= 3 - 4 mice. Data represent the mean value ± SEM. *p< 0.05.

**Figure EV5.**
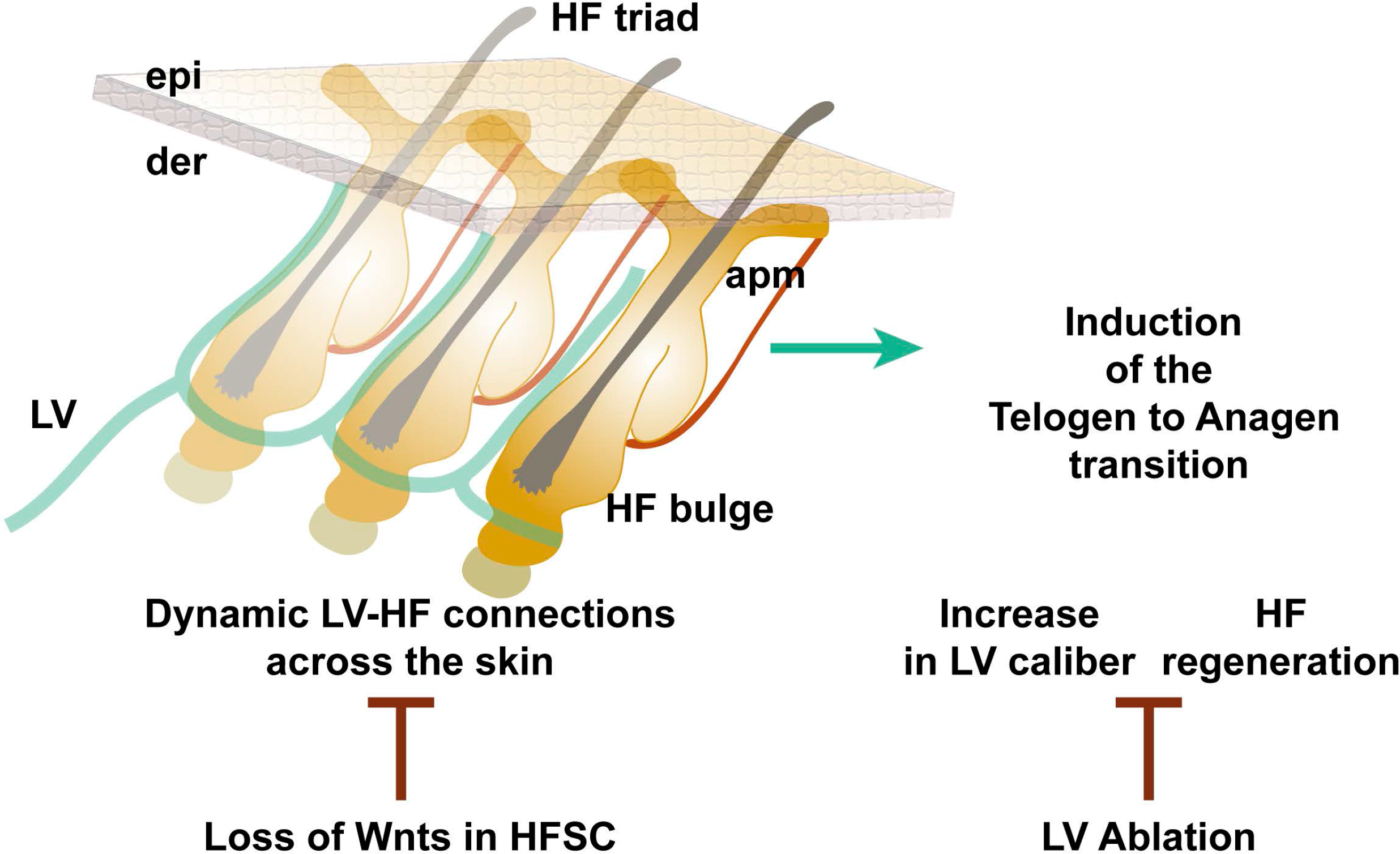
Model of the dynamic interactions of LV with HF in the mouse backskin. LV associate with the posterior permanent side of individual HF. At the level of the HF bulge, LV radiate and converge interconnecting three HF units that in turn, associate with adjacent HF triads across the skin. At the onset of the physiological HFSC activation, or upon pharmacological or genetic induction of HFSC activation, LV expand their caliber suggesting an increased tissue drainage capacity. The loss of Wls ligands in HFSC abrogates the organized association of LV with HF and the ablation of LV blocks the pharmacological induction of HF regeneration. These findings expose LV as components of the HFSC niche, coordinating HF connections at tissue-level and provide insight into their functional contribution to HF regeneration. epi, epidermis; der, dermis; HF, hair follicles; LV, lymphatic vessels; apm, arrector pili muscle.

## Supplementary Materials

Include three supplementary movies, Movie EV1, Movie EV2 and Movie EV3

**Movie EV1. 3D projection of P70 mouse backskin**, using LYVE1 (green) as lymphatic endothelial marker and counterstained with DAPI (blue). 8 fps. Bar, 50 μm.

**Movie EV2. 3D projection of whole mount immunofluorescence analyses of P70 mouse backskin,**using LYVE1 (red) as lymphatic endothelial marker and counterstained with DAPI (blue). 24 fps. Bar, 50 μm.

**Movie EV3. Intravital microscopy in the backskin of the Prox1CreERT2; Rosa-LSL-eYPF mice**. LV flow through triads of HF and across aligned HF rows in the backskin. 8 fps. n= 3 - 4 mice. Bar, 50 μm.

## Appendix.

Includes three tables. Table S1, Table S2 and Table S3.

**Table S1.**
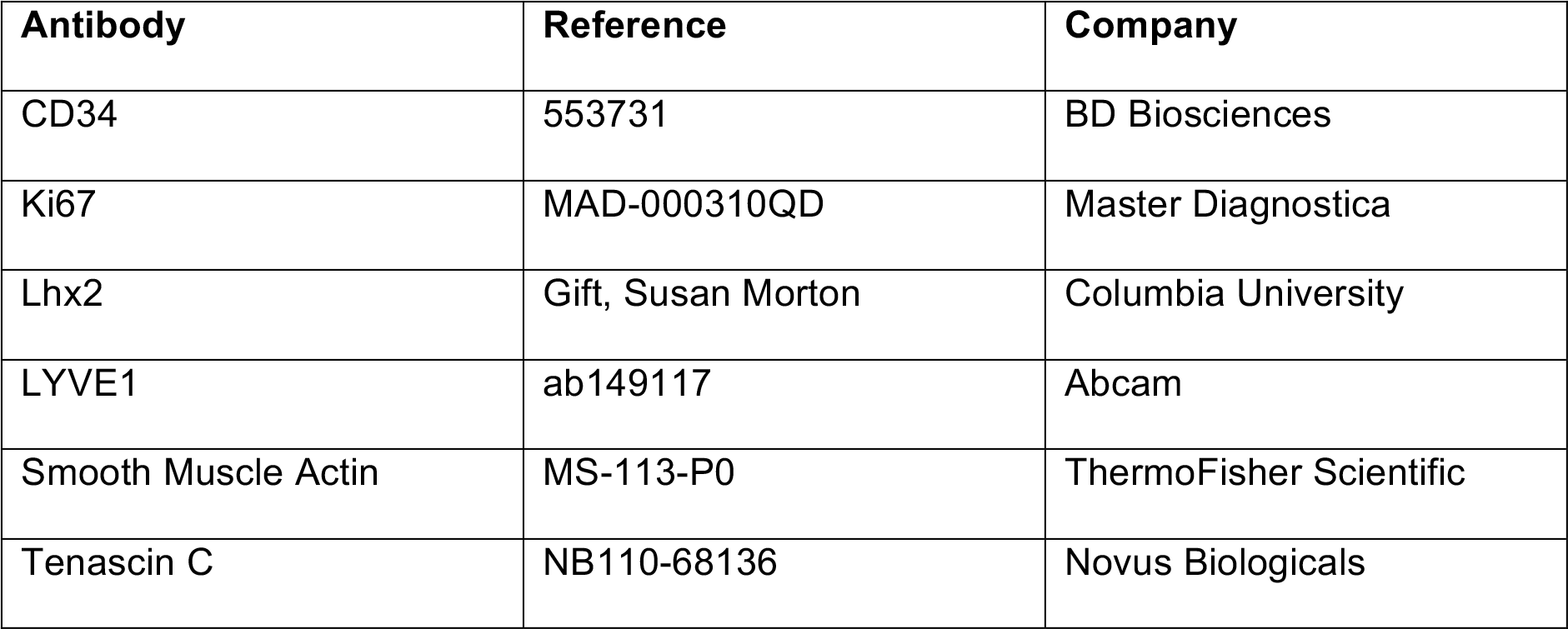
Primary antibodies.

| Antibody | Reference | Company |
| --- | --- | --- |
| CD34 | 553731 | BD Biosciences |
| Ki67 | MAD-000310QD | Master Diagnostica |
| Lhx2 | Gift, Susan Morton | Columbia University |
| LYVE1 | ab149117 | Abcam |
| Smooth Muscle Actin | MS-113-P0 | ThermoFisher Scientific |
| Tenascin C | NB110-68136 | Novus Biologicals |

**Table S2.** Secondary Antibodies.

| Antibody | Reference | Company |
| --- | --- | --- |
| Anti-Rabbit FITC | 711-095-152 | Jackson immunoresearch |
| Anti-Rabbit Alexa Fluor 594 | 711-585-152 | Jackson immunoresearch |
| Anti-Rat Alexa Fluor 594 | 712-585-150 | Jackson immunoresearch |
| Anti-Mouse Alexa Fluor 488 | 715-545-151 | Jackson immunoresearch |

**Table S3.**
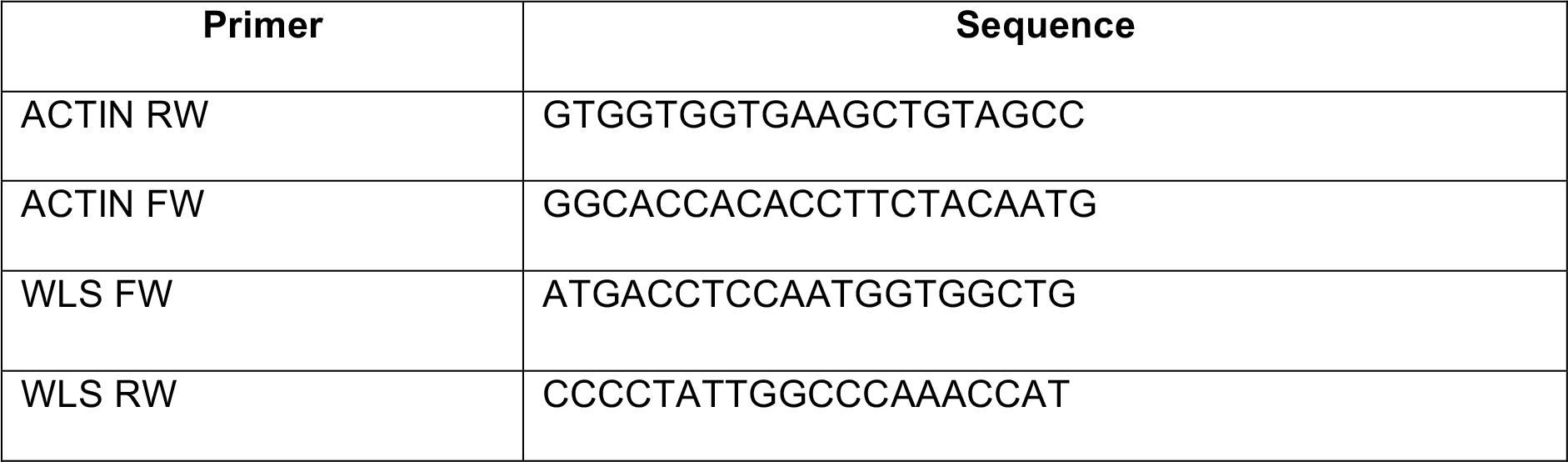
List of RT-qPCR primers.

